# The Crossmodal Congruency Task as a measure of intuitiveness of sensory feedback in the lower limb

**DOI:** 10.64898/2026.08.07.743356

**Authors:** Rohit Bose, Bailey Petersen, Charles Oduro, Roberta Klatzky, Lee Fisher

## Abstract

People with lower limb amputation lack somatosensory feedback from their prosthesis, and this loss contributes to functional deficits, including balance and gait impairments. Recent advances in neuroprostheses have demonstrated that electrical stimulation of sensory nerves in the residual limb and spinal cord can restore lost sensations in the lower limb. To maximize the efficacy of these somatosensory neuroprostheses, the restored sensations should be intuitive, seamlessly integrating into the sensorimotor network. However, it is challenging to quantify the intuitiveness of these evoked sensations. Recent studies have proposed using crossmodal congruency effect (CCE) tasks for this purpose in people with upper-limb amputation. The current study tests the feasibility of the CCE task for assessing the intuitiveness of sensory feedback in the lower limb. We hypothesized that CCE score would reliably differentiate between a more natural (pneumatic) sensation and a less natural (electric) sensation at two locations: the knee and the foot. Across fifteen able-bodied individuals, we observed that the CCE task differentiates sensory modalities at the knee, but not at the foot. Identification of external factors affecting the CCE is needed before it can be implemented to measure intuitiveness of sensory feedback in lower-limb amputees.

## INTRODUCTION

After an amputation, the lack of somatosensory feedback from the prosthesis can lead to substantial functional impairments during balance and gait^1^. Restoring this lost somatosensory feedback may be critical for improving these functional deficits. To this end, a variety of techniques have been demonstrated to restore or augment somatosensory feedback, including sensory substitution^2^ (using a separate modality to compensate for the lack of sensory feedback from the missing limb) or sensory restoration using peripheral nerve stimulation^3,4^ or spinal cord stimulation^5^ to evoke sensations that appear to emanate from the missing limb. With sensory substitution, which often uses visual or auditory cues to replace the lost somatosensation, the sensory feedback is generally not inherently intuitive to the user and a learning period is required to be able to interpret the feedback reliably^6^. This need for interpretation of unintuitive feedback can increase the cognitive load required for maintaining balance and stable gait, hindering performance during tasks where attention is drawn elsewhere, in which case falls are more likely^7,8^. Similarly, during sensory restoration, stimulation of the afferent fibers in the residual limb or spinal cord synchronously activates large and multimodal populations of fibers, often evoking paresthetic sensations in the missing limb^9,10^. Participants in these studies most commonly report experiencing buzzing or tingling sensations, instead of more “natural” sensations, such as touch or pressure. We hypothesize that feedback that is more “natural” and intuitive will be more easily integrated with one’s neural schema, via multisensory integration, with the potential to provide greater functional benefits to the user.

Recent studies have investigated the use of biomimetic stimulus patterns that more closely match the patterns of natural activity in somatosensory afferents. Seminal work in individuals with upper-limb amputation have shown enhanced improvements in task performance when using these biomimetic patterns^11–13^. However, we do not yet know whether the subjective experience and intuitiveness of these sensations play a role in the enhanced task performance, largely due to a lack of reliable metrics that can quantify multisensory integration of sensory feedback. Alternatively, it is possible that the improved task performance occurs because biomimetic stimuli can better engage with spinal and cerebellar circuits, regardless of the perceptual qualities of the evoked sensations. To date, research on somatosensory neuroprostheses has relied heavily on participants’ subjective “naturalness” ratings, which vary in definition and are often highly variable both within and across participants^13,14^. Additionally, “naturalness” ratings do not assess how intuitive a sensation is to the user. The intuitiveness of feedback from a sensory channel lies in its integration with the multiple perceptual pathways that govern motor performance, especially vision^15,16^. Thus, a measure of multisensory integration is necessary to evaluate the intuitiveness of stimulation in sensory neuroprostheses.

The crossmodal congruency effect (CCE) task was introduced to assess the interaction of different sensory modalities in the body, and thus can serve as an assessment of multisensory integration^17,18^. The participant is instructed to report the location of a target stimulus delivered in one modality (e.g., electrical stimulation) while ignoring a simultaneous distractor stimulus in another modality (e.g., visual), appearing at a similar or different location. With this paradigm, participants’ reaction times are typically slower in reporting the location of the target sensation when the distractor is at a different location on the body (i.e., an incongruent trial) as compared to when the target and distractor are at the same location (i.e., a congruent trial). This disadvantage in response times for incongruent trials compared to congruent trials (called the crossmodal congruency effect; CCE) is assumed to occur because a remote distractor is encoded as a discrete event that competes more effectively with the target than a co-located distractor. Thus the CCE score serves as a measure of multisensory integration of signals at the target location, with a higher score indicative of greater integration. Along with behavioral measures, past studies have also used neural signatures from electroencephalography (EEG) to quantify multisensory integration and incongruency effects^19–21^. The incongruency effect has been shown to increase the negative (N2) component of event related potentials^19^, along with an increase in the theta and gamma band power^18,20^ in the central and parietal regions.

Blustein, et al., proposed the use of CCE scores to evaluate effectiveness of different types of artificial sensory feedback for use in neuroprosthetic studies^17^. They found that it was more challenging for users to ignore a visual distractor and attend to a mechano-tactile stimulus than an electrical stimulus, indicating greater integration of vision with tactile than with electrical inputs. Their work suggested the use of CCE scores as a proxy for intuitiveness of sensory feedback in neuroprosthesis studies for individuals with upper-limb amputation. Another study used the CCE task to quantify embodiment of an upper-limb closed-loop prosthesis^22^. Both of these studies focused on quantifying somatosensory feedback in the upper limb, though the field of somatosensory neuroprosthetics has recently expanded to target individuals with lower-limb amputation. As such, an adapted version of this task should be validated for the lower extremities. This study aimed to develop a crossmodal congruency task to assess the multisensory integration of somatosensory restoration for people with lower-limb amputation. As a first step, our goal was to validate this measure in able-bodied participants, with the future goal of using CCE scores as a measure of multisensory integration of sensory percepts in neuroprosthetic studies.

## RESULTS

We performed experiments in fifteen able-bodied individuals to validate our proposed CCE paradigm as a measure of intuitiveness of sensory feedback for lower-limb somatosensory neuroprostheses. Our approach was built on the assumption that a pneumatic tactile stimulus would be more intuitive than an electrical stimulus, reflecting the neural populations these channels activate. Specifically, while a pneumatic tactile stimulus recruits slowly adapting type 1 fibers (which respond to skin indentation), as well as rapidly adapting and PC fibers (responding to change in pressure) in a finely tuned encoding of signals, electrical stimulation recruits large populations of afferent fibers simultaneously. The differences in neural populations recruited by these stimuli suggest that pneumatic and electrical stimuli should have distinct CCE scores and naturalness ratings. Thus, we hypothesized that the CCE score would be higher for pneumatic stimuli than electrical stimuli. We delivered both stimuli at the knee and the foot because recent studies from our lab and others have reported that both peripheral nerve and spinal cord stimulation evoke sensations in these locations in people with lower-limb amputation^5,23,24^. However, the perceived visual distance between the knee and the foot is larger than the distances reported in previous CCE studies^17,25,26^. As such, the perceived brightness of the visual stimulus could also differ between the knee and foot location due to the variable distance from the eye and differences in ambient light. Since stimulus intensity can affect the CCE^27^, we first confirmed whether the visual stimulus is perceived with similar time delay from the two locations using a visual reaction time test. Subsequently, and prior to measuring CCE scores, we also matched the electrical and pneumatic stimulus intensity for each location as described below.

### Visual Reaction Time Test

To measure visual reaction time, a visual stimulus was delivered using LEDs with three different luminous intensities (33%, 66% or 100% of maximum brightness). Participants performed a speeded-response task where they verbally report the location of the visual stimulus as quickly as possible. We used verbal responses instead of button presses to eliminate differences based on handedness and the potential variations in response time with a lateralized stimulus^28,29^. In 12 out of 15 participants, we did not observe any significant differences in reaction times across the two locations and luminous intensities (see Supplementary Table 1, Supplementary Fig 1). For the other three participants, post-hoc analysis further revealed significant differences only between the lower luminous intensities (33% and 66%). Therefore, we selected the maximum intensity of the visual stimulus for the CCE experiment.

### Stimulation Amplitude

Another factor that can affect the CCE scores is stimulus intensity. Therefore, to match the perceived intensity between the pneumatic and electrical sensations, we varied the amplitude of the electrical stimulus while keeping the pneumatic stimulus intensity constant. The pneumatic and electric stimuli were delivered as a pair, and the participant was instructed to answer which stimulus felt stronger. Based on the response, we varied the electric stimulus until the participant responded that both stimuli were perceived with equal intensity. We repeated this process separately for the knee and the foot. Overall, we observed that the thresholds were higher at the foot than the knee, and therefore, electric stimulus amplitudes delivered to match the pneumatic stimulus were consistently higher in the foot compared to the knee (Wilcoxon’s signed rank test, p<0.0001).

### Crossmodal Congruency Effect Task

The crossmodal congruency effect (CCE) task used a somatosensory stimulus (i.e., the target) simultaneously delivered with a visual stimulus (i.e., the distractor) (Fig 1). The target and the distractor were delivered at a spatially congruent location (i.e., both at knee or foot) or incongruent locations (i.e., target at the knee and distractor at the foot or vice-versa). The primary outcome, CCE_RT_, was the difference in the reaction time to the tactile stimulus for the incongruent condition and the congruent condition. We also calculated the number of incorrect responses during the congruent and incongruent conditions. The difference between the two is referred to as CCE_Error_. Previous CCE studies have performed statistical analysis at the group level. However, for neuroprosthetic applications, perception of the artificially evoked sensation varies between individuals. Therefore, to effectively measure intuitiveness of sensory feedback, we believe it is imperative to evaluate effects on an individual basis. Hence, we performed our statistical analyses to measure differences within individual participants (i.e., with paired statistics).

**Fig 1:**
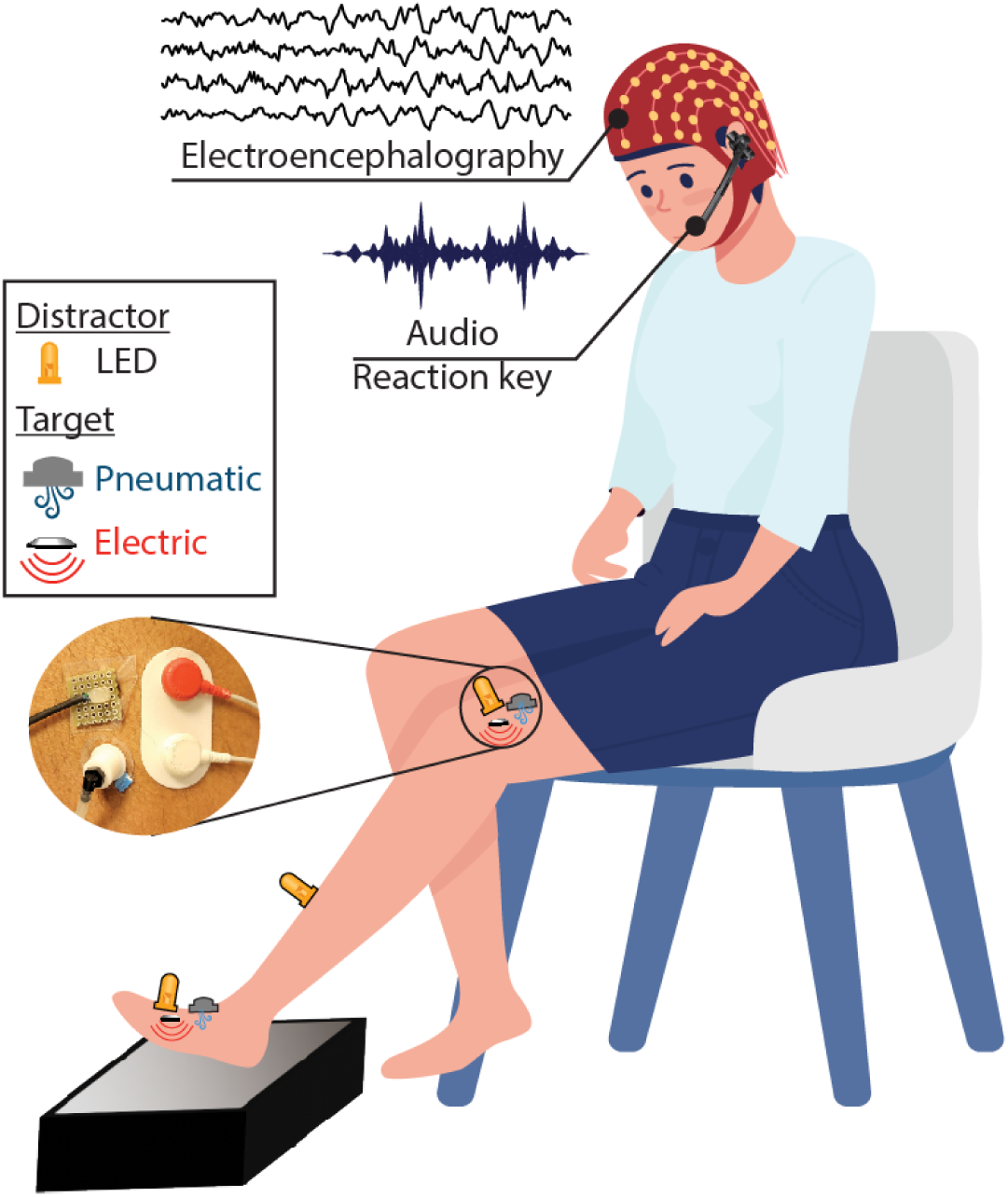
Experimental Setup. The participants were seated and received target tactile (pneumatic and electric) stimulation along with a distractor visual stimulus at the knee or foot. Verbal responses and EEG signals were recorded during each trial.

Reaction times for the incongruent trials were higher overall than for congruent trials for both stimulus modalities and locations for all participants, except one subject for the pneumatic-knee stimulus and another subject for the electric-foot stimulus (see Supplementary Fig 2). However, significant CCE scores (differences between the incongruent and congruent trials) were only present for 67% of the participants. Out of 15 participants, we observed significant differences in 10 participants for pneumatic-knee stimuli, 11 participants for electric-knee stimuli, 9 participants for pneumatic-foot stimuli, and 10 participants for electric-foot stimuli. However, among the participants with significant differences, the result was not consistent for both stimulus modalities. Only 9 and 8 participants had significant CCE scores for pneumatic vs electric stimuli at the knee and the foot, respectively.

We computed the CCE scores for participants with significant differences between congruent and incongruent trials for both stimulus modalities and compared scores at the knee and the foot (Fig 2). At the knee, 14 of 15 participants had a higher CCE_RT_ for pneumatic stimuli compared to electric. At the foot, only 3 participants had a higher CCE_RT_ for pneumatic stimuli as compared to electric, whereas 5 participants had the opposite trend (Fig 2).

**Fig 2:**
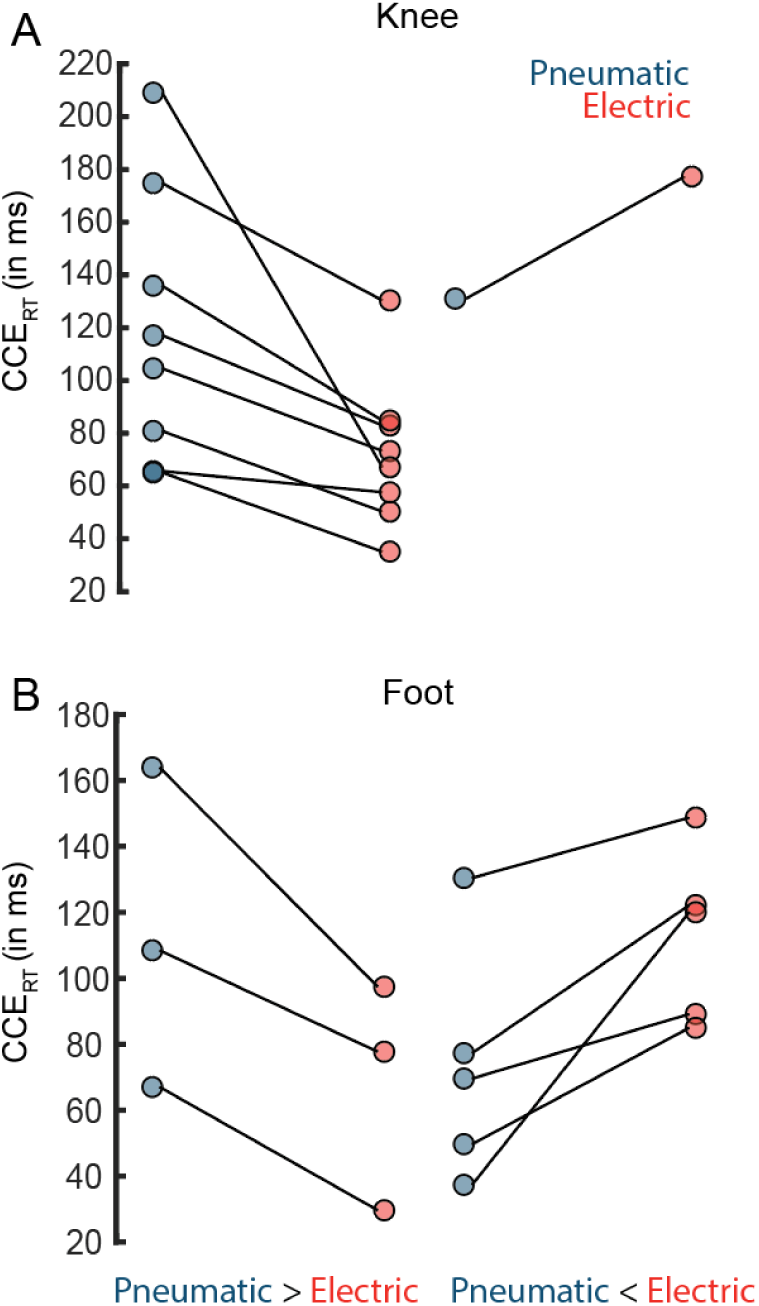
CCE_RT_ for all participants at the (A) knee and (B) foot. The left paired plots represent the participants with a higher CCE_RT_ for pneumatic compared to electrical stimulus and the right paired plots represent the participants with a higher CCE_RT_ for electric compared to pneumatic stimulus.

Overall, the incorrect responses constituted fewer than 9% of the total trials. Similarly to CCE_RT_, we observed relatively consistent results at the knee, but not at the foot. Ten participants had higher CCE_Error_ for the pneumatic stimulus at the knee, while only 4 participants had higher CCE_Error_ for the pneumatic stimulus at the foot. We did not observe any consistent patterns across locations or stimulus type for the CCE_Error_.

### Behavioral Measures

Fatigue often plays a role in psychophysical experiments. To minimize fatigue, the entire experimental session was divided into six sets. At the end of each set, participants rated their fatigue level on a 0-10 Likert scale. All participants reported slightly higher fatigue scores at the end of the experiment (difference between the last and first session, mean ± standard deviation, 1.47±1.36). However, we did not observe any significant correlation of the fatigue scores and the CCE_RT_ for pneumatic-knee (r=-0.3, p=0.29), pneumatic-foot (r=-0.01, p=0.96), electric-knee (r=- 0.15, p=0.58) or electric-foot (r=0.21, p=0.46) conditions.

Though not a primary outcome of our study, participants also reported their perceived naturalness for the electrical and pneumatic stimulus modalities at each location on a 0-10 Likert scale (see Supplementary Fig 3). 8 and 10 participants had a significant difference for pneumatic vs electric at the knee and foot respectively (tested via bootstrapping; see Methods for additional information). We also evaluated the relationship between the difference in naturalness ratings and CCE score differences at the knee and the foot (Supplementary Fig 4) across all participants. We did not observe a significant inter-participant correlation (Pearson’s r) between the CCE scores and the perceived naturalness ratings for pneumatic-knee (r=-0.03, p=0.91), pneumatic-foot (r=-0.17, p=0.54), electric-knee (r=-0.20, p=0.47) or electric-foot (r=0.04, p=0.9) conditions. These results suggest that the CCE_RT_ score is not a proxy measure for a perceived naturalness rating.

### Reaction Time Variability

We observed variability in reaction times across participants (interquartile range of congruent trials: 439-1141 ms; incongruent trials: 404-1176 ms). Such variability might affect the CCE_RT_ as delayed reaction times may reduce the CCE. However, we did not observe any significant correlation (Pearson’s correlation) between congruent reaction times and the CCE_RT_ for pneumatic-knee (r=0.25, p=0.3), electric-knee (r=0.06, p=0.85), pneumatic-foot (0.29, p=0.30) or electric-foot (r=-0.02, p=0.95) conditions.

### Group-Level Analysis

Previous studies using the CCE paradigm to assess upper-limb somatosensory neuroprostheses performed group-level analysis, in which they obtain a single CCE score for each participant and then perform statistical analysis combining all participants^17,18,30^. To compare our findings with those studies, we performed similar group-level statistical analyses (Fig 3). With this approach, there were significant differences between the incongruent and congruent trials for all conditions (pneumatic-knee: p<0.005; electric-knee: p<0.0001; pneumatic-foot: p<0.0001; electric-foot: p<0.0001). At the knee, CCE_RT_ was higher for the pneumatic stimulus (91.15 ± 77.1 ms) compared to the electric stimulus (74.36 ± 47.37 ms). At the foot, CCE_RT_ was higher for the electric stimulus (81.08 ± 43.24 ms) compared to the pneumatic stimulus (77.64 ± 33.40 ms). The difference across stimulus modalities was greater at the knee compared to the foot.

**Fig 3:**
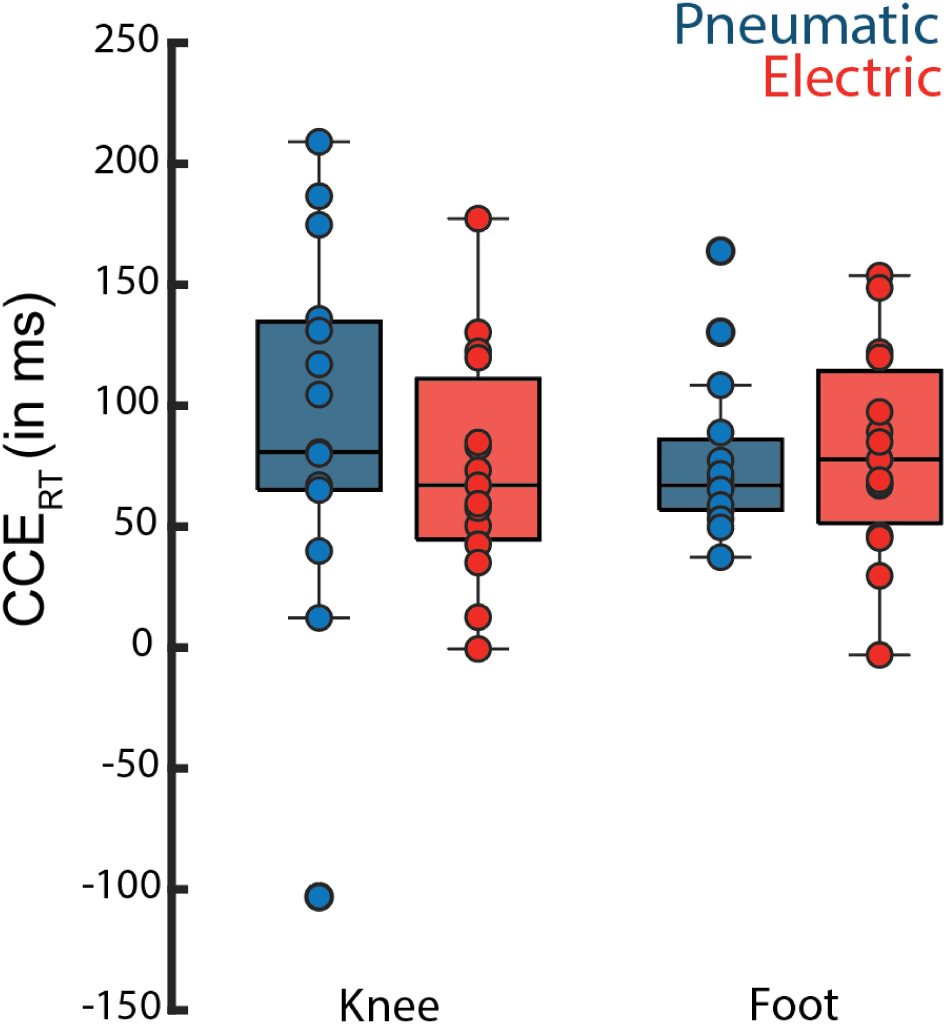
Group-level CCERT for Pneumatic and Electric stimulus at the knee and foot location.

### EEG Band Power

We collected EEG signals from 12 out of 15 participants to find cortical biomarkers of intuitiveness^19–21^. We analyzed the band power of the EEG signal during the 200 msec interval after stimulus delivery. Overall, we found the delta (0.3-4 Hz) and theta (4-8 Hz) bands to be the most relevant features to differentiate congruent and incongruent conditions. The difference was primarily observed in the parietal region (Fig 4,5; Supplementary Fig 5-7). We observed variability across the knee and foot location. At the knee, incongruent trials had higher parietal power for both delta and theta bands compared to the congruent trials (Fig 4,5). At the foot, we observed the opposite patterns, where the congruent trials had higher parietal power compared to the incongruent trials (Fig 6), though the difference between conditions was not statistically significant for all participants and the pattern of significant differences vs. location and modality was highly variable. Out of the 12 participants, two had a significant difference in parietal power for both stimulus modalities at the foot (Fig 5). At the knee location, four participants had a significant difference in parietal power for the electric stimulus but not for the pneumatic stimulus (Fig 4). We also did not observe any significant difference in the group-level analysis for any electrode or frequency band (Supplementary Fig 7).

**Fig 4:**
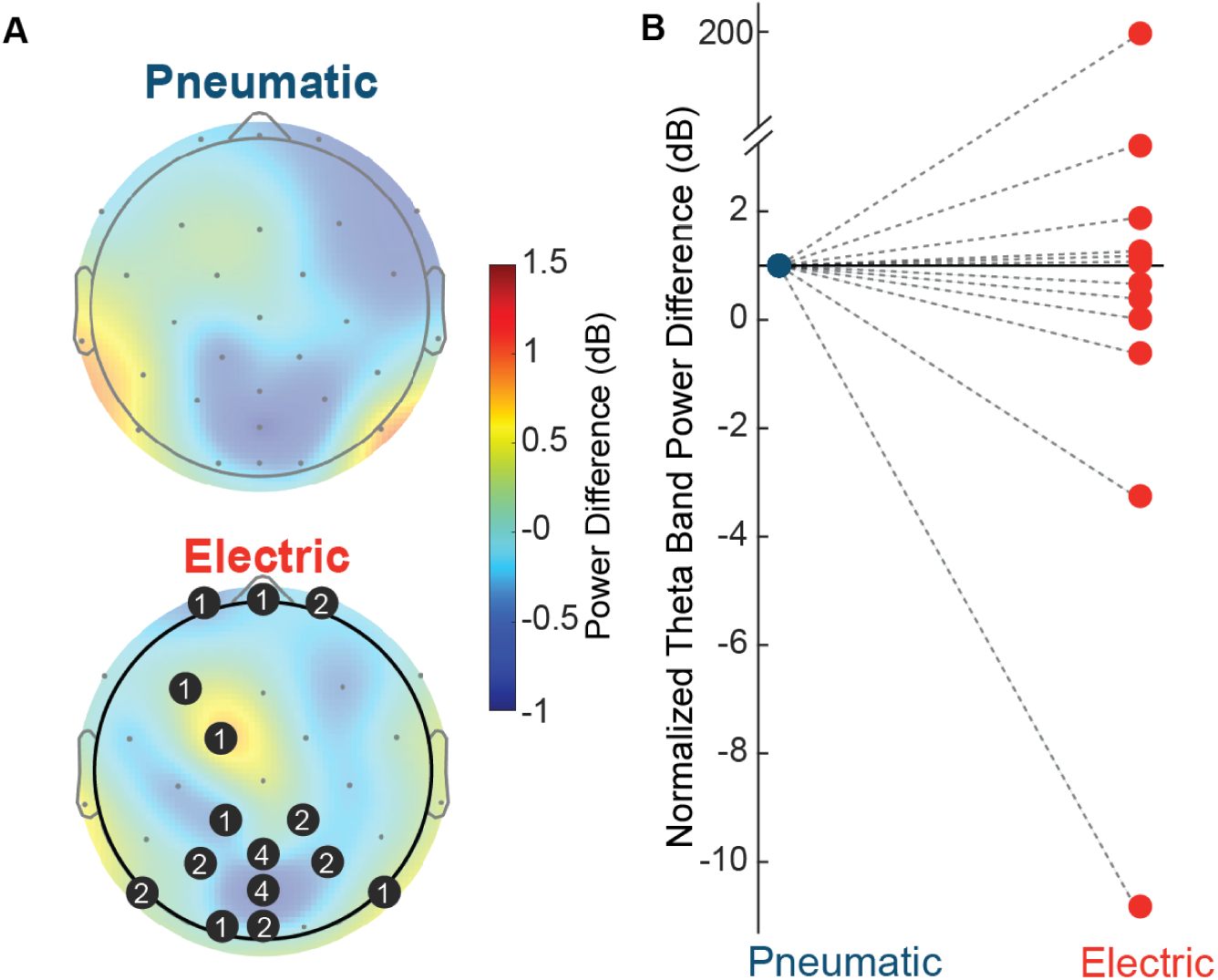
Stim-locked theta band power results for stimulation at the knee. (A) Topographic plots show the difference between the incongruent and congruent conditions for the pneumatic and electric stimuli. The black circles indicate the number of participants for which the electrode was significantly different. (B) Difference in the theta band power between the incongruent and congruent conditions for the electric stimulus normalized to the pneumatic stimulus for each participant. The dashed lines indicate that the incongruent theta band power was not significantly different from the congruent theta band power for pneumatic and electric stimuli.

**Fig 5:**
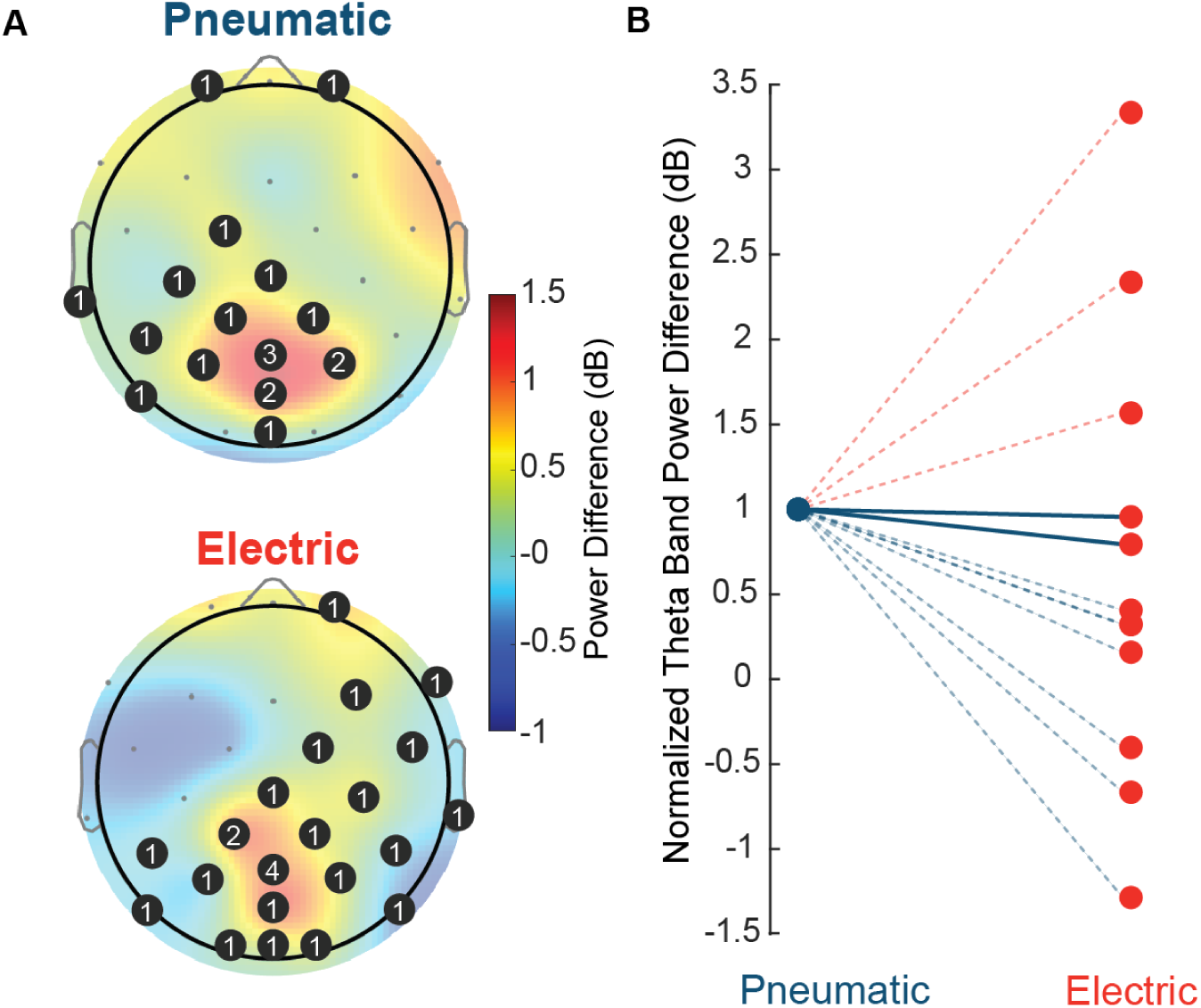
Stim-locked theta band power results for stimulation at the foot. (A) Topographic plots show the difference in the incongruent and congruent conditions for the pneumatic and electric stimuli. The black circles indicate the number of participants for which the electrode was significantly different. (B) Difference in the theta band power between the incongruent and congruent condition for the electric stimulus normalized to the pneumatic stimulus for each participant. The solid lines indicate that the incongruent theta band power was significantly different from the congruent theta band power for both pneumatic and electric stimulus and dashed lines indicate that there was no significant difference.

**Fig 6:**
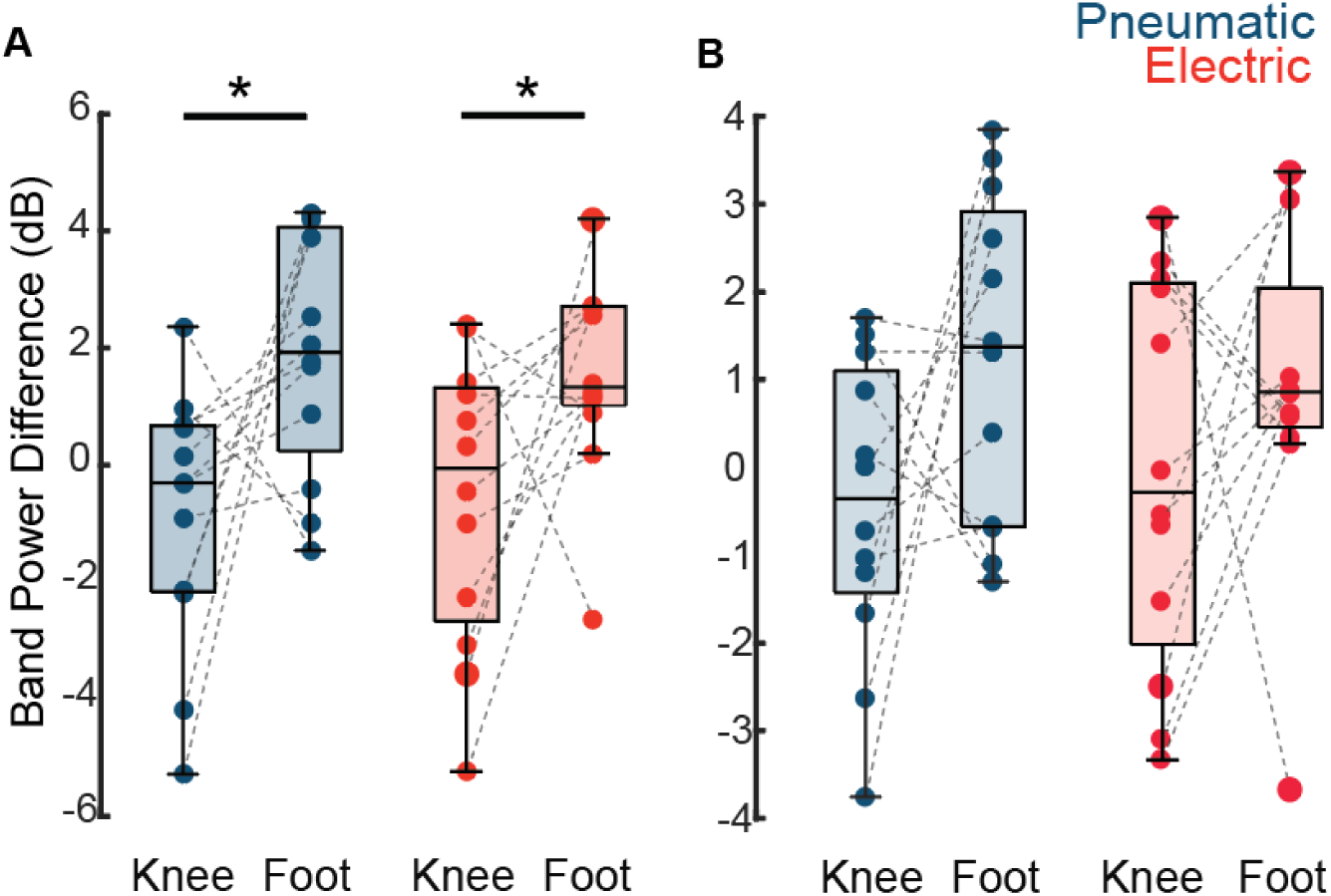
Comparison of band power difference between incongruent and congruent condition for (A) delta and (b) theta band power between the knee and foot location. * indicates a significant difference between the knee and foot condition (Wilcoxon rank sum test, p<0.05).

## DISCUSSION

In this study, our goal was to adapt the existing CCE task paradigm to quantify ‘intuitiveness’ of sensory feedback for people with lower-limb amputation. We assumed that a pneumatic stimulus would be more intuitive than an electrical stimulus based on differences in afferent fiber recruitment and hypothesized that it would therefore have a higher CCE_RT_. However, the congruency effect was present in 60% of the participants at the knee and 53% at the foot. Within those subjects, we observed consistently higher CCE scores for the pneumatic stimulus at the knee, whereas results at the foot were less consistent. The lack of correlation between CCE scores and ratings of naturalness further indicates that, at least for the lower limb, CCE is not simply a proxy for subjective naturalness. Below we further discuss potential explanations for these results and limitations of the CCE task for somatosensory neuroprostheses.

In past studies that used CCE tasks to quantify evoked sensations in the upper limb, researchers have selected the thumb and index finger of the same hand^17^, tip and base of the index finger^31^, two adjacent fingers^25^ or two fingers in either hand^32^. Other studies have shown that the CCE is lower if the target-distractor pair is on different sides of the body compared to on the same side^33,34^. Blustein, et al., demonstrated the effect of relative distance between the target and stimulus within a congruent location^17^. To date, none of the studies have investigated the effect of the distance between the incongruent target and the distractor locations on the same side of the body. Because spinal cord stimulation^5^ and peripheral nerve stimulation^23,24^ have been shown to evoke sensations that emanate from regions of the knee and foot in people with lower limb amputations, we selected these locations in our experimental implementation of the CCE task. The knee and the foot are spatially further from each other than the locations used in prior studies of the upper limb, which may reduce the CCE.

The quality of evoked sensations plays a major role in the perceived naturalness of the sensations^35^. Though the majority of participants (8 for the knee and 10 for the foot) had significant differences in naturalness ratings between the two modalities, not all participants rated the pneumatic stimuli as more natural. Additionally, the overall similarity in subjective naturalness ratings was surprising. Anecdotally, some subjects reported that the perceived quality of the two modalities at the foot were similar, which may also have led to inconsistent results in the foot. Despite a lack of relationship between naturalness ratings and CCE scores, other types of sensations that may be more clearly distinct from electrical stimulation may be useful in future studies.

Tactile sensations are conveyed primarily via Aβ afferent fibers. These fibers are not homogenously distributed across the body, which leads to different levels of spatial acuity in different regions of the body. Previous studies primarily stimulated the fingers which have the highest density of receptors and highest spatial acuity^30,36,37^. The lower extremity has lower spatial acuity than the fingers, potentially limiting direct comparisons with prior studies in the upper extremity. Within our study, the two locations differ in spatial acuity, as well. The foot has greater spatial acuity^36^ compared to the knee. Further research is required to understand the effect of stimulus location and spatial acuity on CCE for its application in sensory restoration studies.

CCE studies have long been analyzed at a group level to understand the integration of multiple sensory modalities among a population^18^. From a cognitive neuroscience standpoint, group-level analysis is meaningful, but a similar approach cannot be adopted for quantifying the ‘intuitiveness’ of sensory feedback from a somatosensory neuroprosthesis. For these applications, we need a measure of intuitiveness for individual users^5,23,24^. Towards that goal, we performed a paired statistical analysis in this study. Our results show that CCE is not present for all participants, either in terms of reaction time or EEG band power measures. Furthermore, we did not find a relationship between CCE scores and subjective naturalness ratings. A recent study used the CCE paradigm to quantify prosthetic ownership with combined sensory feedback of touch and kinesthesia at the participant level^22^, and observed the expected result in only one out of two participants. These findings question whether the CCE paradigm is an appropriate measure of ‘intuitiveness’ of sensory feedback at the individual level.

Our findings could also be affected by other factors that impact CCE scores. In the field of cognitive neuroscience, there is ongoing debate about the relative contributions of multisensory integration, attentional demands, and response conflict on CCE scores^18,34^. Response conflict is an incorrect mental representation of sensation locations that involuntarily occurs when a distractor stimulus appears at the same time^34^. A study evaluating these relative contributions determined that response conflict and, to a lesser extent multisensory integration, were the primary mediators of the standard crossmodal congruency task^34^. In the traditional crossmodal congruency task, a block is held with the index finger and the thumb in each hand with LEDs and tactile stimulators at each location^32^. Participants are asked to discriminate between location (index finger or thumb) while visual distractors appear on either hand (same or opposite hand of the stimulus) on the same finger (congruent) or the other finger (incongruent). The slowing of response times in incongruent trials represents the amount of time it takes to inhibit these incorrect representations of the stimulus, regardless of the location of the distractor (same or opposite hand incongruent trials). However, multisensory integration is location dependent and can thus explain the increase in CCE score on only the ipsilateral hand. If only response conflict were at play, the CCE score would be similar on both hands, as the location of the distractor should not matter, but in those experiments, the CCE score is higher on the ipsilateral hand than the contralateral hand. Thus, both response conflict and multisensory integration have a role in the crossmodal congruency task.

Response conflict can be considered an indicator of the interference or dominance of visual stimuli on somatosensory stimuli, while multisensory integration is an indicator of how well the visual and somatosensory stimuli merge in the neural schema. Both aspects of crossmodal perception are important, however for the purposes of evaluating varying types of somatosensory stimuli, assessing multisensory integration in lieu of response conflict may be more appropriate. To eliminate the contribution of response conflict, a shift to a “Go-No Go” task would be necessary. In this task, the participant is asked simply to report if they felt a sensation, not to identify its location.

There is also a discrepancy in how past studies have recorded participant responses. Depending on the task, studies have used button presses with fingers^25,31^, pedal presses with both feet^17,38^ or pedal release with the heel and the toe^32–34,39–41^. The reaction time for these methods can differ based on the musculature involved. Also, bilateral reaction times differ from unilateral reaction times. Past studies have shown that vocal responses are more robust to age-related differences than manual button press reaction times^42^. Therefore, we selected vocal responses to measure reaction time. We instructed participants to add the phoneme ‘t’ before saying ‘knee’ or ‘foot’ to avoid unwanted discrepancy in our ability to detect reaction time from the recorded auditory signals. Notably, vocal responses are slower than a button press^41^, which would affect the reaction times in comparison to previous studies, but not the CCE scores themselves.

Some studies have shown that EEG-based cortical biomarkers are associated with the CCE^19–21^. However, we did not observe a consistent pattern among the EEG features at an individual level. The lower frequency bands in the parietal region are most relevant for conflict processing and crossmodal integration^19,43–45^. The delta band (0.3-4 Hz) is primarily involved in cognitive functions like attention, working memory and response inhibition^46^. The theta band (4-8 Hz) is primarily responsible for stimulus processing and multimodal sensory integration^47^. In the congruent conditions, when the visual and somatosensory stimulus are in the same spatial location, multimodal sensory integration should be stronger, which has been observed in prior studies showing higher band power compared to the incongruent conditions^19^. We observed a similar effect in the knee location, but an opposite effect at the foot. This may be because the CCE did not occur at the foot. However, the lack of a consistent pattern in the EEG band power features between the two stimulus modalities at both locations further calls into question the appropriateness of the CCE task as a measure of intuitiveness at the individual level.

A major limitation of this study is that we tested the CCE paradigm as a measure of naturalness in able-bodied participants with electric and pneumatic stimulation. People with amputations and lost sensations might have higher embodiment of the artificial sensation and hence could have a different CCE score. Another limitation of this study was that we were unable to match the perceived intensity of the pneumatic stimulus at the knee and the foot. Because higher intensity stimuli are more salient and elicit faster reaction times, the CCE score for higher intensity stimuli would decrease, as the distractor is less effective at overcoming the target stimulus effect. Additionally, some participants reported numbness in the distal foot with their leg extended resting on a step for 2-3 hours. Although we took longer breaks in these cases to allow the participants to relax their legs, the effect of numbness within each session could not be controlled and could influence CCE scores. In future studies, reducing the experiment time could ameliorate this effect. Further, longer experiment durations can induce learning effects in the CCE score^17^.

## CONCLUSION

In summary, while the CCE score was an indicator of multisensory integration of stimuli in the knee, this finding did not generalize to the foot. Furthermore, differences in CCE scores were not correlated with differences in subjective reports of naturalness for the stimulus modalities. Given these results, additional steps need to be taken to further develop a more useful proxy of naturalness and a more appropriate method of measuring multisensory integration of stimuli and evaluating the functional effects of somatosensory neuroprostheses.

## METHODS

### Participants

Twenty able-bodied individuals were recruited for the study, and fifteen individuals completed the entire study. Inclusion criteria were adults between the ages of 18 and 65 who had normal or corrected to normal vision. Participants were excluded if they: (1) had a history of neurological disease, motor impairment, or chronic pain, (2) had an implanted electrical device, port, or pump, (3) were being treated for cancer or were in acute remission, (4) were pregnant, or (5) had any implanted metal hardware anywhere in their body. The experimental protocol was approved by the University of Pittsburgh’s Institutional Review Board.

### Experimental Setup

Participants were seated in a chair and resting their right heel on an elevated platform (20 cm) as shown in Fig 1. The two somatosensory stimuli (target) and the LED stimulus (distractor) were placed on the mid-thigh (referred to as ‘knee’, 10 mm proximal to the superior patella) and the dorsal foot (referred to as ‘foot’, 5 mm distal to the midpoint of the two malleoli).

The somatosensory stimulus was delivered using the Galileo tactile stimulation system (Epic Medical Concepts & Innovations, Inc., Mission, Kansas USA). The stimulus was delivered through a 26-foot-long polyurethane tube. Polymer based stimulator probes were attached to the skin using a double-sided tape to fix the location of the stimulus delivery. The pneumatic stimulator was placed in a different room so that sound from the device did not bias the participants.

The electrical stimulus was delivered using a DS8R Stimulator (Digitimer Ltd., England, UK). Bi- phasic charge balanced stimulation trains with 200 µs pulse width and 50Hz frequency for 1 second were delivered through bipolar pairs of surface electrodes (Ag/AgCl disposable dual electrodes, MVAP Medical, Thousand Oaks, CA) attached to the skin.

We also placed two white light-emitting diodes (LEDs) at each of the locations. Additionally, there was a fixation LED placed midway between the foot and knee LED on the tibia. An Arduino board was used to trigger the LEDs.

Participants were asked during the experiments to respond verbally to evoked stimuli with “t-knee” or “t-foot,” based on the location of the stimulus. Verbal reaction times were recorded with a Cedrus Voice Reaction device (SV-1 Voice Key).

### Visual Reaction Times

To ensure the longer visual distance to the foot LED was not affecting reaction times, we first recorded reaction times in response to visual feedback only. Participants were asked to fixate on the center LED and then respond with “t-knee” or “t-foot,” based on the location of the visual stimulus. Verbal responses were analyzed to determine incorrect responses, which were removed prior to analysis.

### Intensity Matching Task

To match the intensity of the electrical and pneumatic stimuli, stimulation was controlled using custom software in MATLAB and both stimuli were presented in randomized order with a 1 sec interval between. The participants were asked to report which stimulus felt more intense. Since amplitude of the pneumatic tactor stimulation was fixed, the electrical amplitude was then increased or decreased by 0.5 mA based on their response. This was repeated until the participant responded in three consecutive trials that the two stimuli were perceived with equal intensity. Intensity matching was completed for the foot and knee electrodes separately and the matched stimulus amplitudes were then held constant throughout the remainder of experiments.

### Stimulus and Distractor Synchronization

The LED distractors were triggered through an Arduino device whereas the pneumatic and electric stimuli were triggered through digital pulses from a Ripple Nomad data acquisition system. The pneumatic stimulus was delivered as a pulse and had inherent delays that were both internal to the system as well as resulting from the inertia of air within the length of the cables. To ensure that the target stimulus and the distractor stimulus were synchronized, we evaluated the hardware lag between the target and distractor stimulus triggers. We used a digital input/output headstage (Ripple Neuro, UT, USA) to measure the timing of the LED along with the triggers for pneumatic and electrical stimuli. We used the Galileo software to determine the cable related delays of the pneumatic stimulus to the participant. Between the LED and the electrical stimulus, the delay was less than 1ms (∼0.1-0.9 ms) for both knee and foot. The internal delay of the Galileo to generate the pneumatic stimulus was 31 ms and the delay for the 26-foot cables was an additional 35 ms. All the measured hardware lags were then adjusted during the stimulus delivery to synchronize the target and distractor stimuli to <1ms delay.

### Crossmodal Congruency Task

In the crossmodal congruency task, somatosensory (target) and visual (distractor) stimuli were delivered simultaneously. The visual stimulus (LED) appeared either in the same location as the somatosensory stimulus (congruent trials) or in the other location (incongruent trials). The participants responded as quickly as possible, stating where they felt the somatosensory stimulus (“t-knee” or “t-foot”) while avoiding the distractor visual stimulus. The phoneme “t” was added preceding the location in verbal responses to avoid the variations in detection delays based on the audio signatures of the words “knee” and “foot”. Participants were instructed to fixate on the center LED throughout the trial. At the beginning of each trial, the fixation LED turned on for 2 s, followed by the target and distractor stimuli. Participants wore headphones with white noise to mask out any sound of the pneumatic stimulation.

Participants completed a total of six sets with 12 blocks each and 5-minute rest breaks between sessions. Each block consisted of randomly assigned incongruent and congruent trials for each stimulus modality (pneumatic or electrical) at each location (knee and foot), with a total of 8 trials per block. Additionally, 6 catch trials per block were added, in which both LEDs were illuminated, and the participant was instructed to respond “invalid.” These trials ensured the participant was attending to the fixation light. Prior to the first session, practice trials were performed until the participant was familiarized with the task. Upon the end of each session, participants reported the perceived “naturalness” of the pneumatic and electrical sensations on a scale of 0-10 (10 being completely natural). Additionally, participants reported their fatigue level on a 0-10 scale (10 being most fatigued) between each session. Incorrect trials and trials with early (<200 ms) or delayed responses (>2000 ms) were discarded prior to analysis (the range was determined from preliminary pilot experiments).

### EEG Data Acquisition and Analysis

EEG data were acquired from 12 participants using Ag/AgCl electrodes according to the international 10-20 standard^48^. The electrodes were placed at 32 locations across the scalp, and the data was sampled at 2Khz (SAGA, Twente Medical Systems International (TMSi) B.V., the Netherlands). Electrode impedance was maintained below 20 kΩ. Pre-processing of the data was performed using the EEGLAB toolbox in MATLAB 2022a. Data were downsampled to 500Hz and band-pass filtered from 0.3-40 Hz and notch filtered at 60Hz to eliminate any power line noise. The filtered data were epoched from 2 sec before stimulus delivery until the end of the trial (marked by the verbal response). Data corresponding to incorrect trials and trials with early (<200 ms) or delayed (>2000 ms) response was not included for further analysis. Independent Component Analysis was used to remove ocular and muscle artifacts^49^.

### EEG Analysis

After pre-processing, each trial was further epoched from stimulus onset to 200 msec post-stimulus onset^19,20^. We calculated the spectral power using a short-time Fourier transform. After that, we calculated the band power for five frequency bands: Delta (0.3-4 Hz), Theta (4-8 Hz), Alpha (8-13 Hz), Beta (13-30 Hz) and Gamma (30-40 Hz), using the following formula^50^:

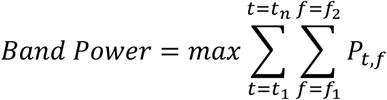

Where *P_t_*_,*f*_denoted the spectral power at time point *t* and frequency *f*, *t*_1_ and *t_n_* denotes the boundary time points and *f*_1_ and *f*_2_ denote the start and end frequency of the respective bands.

### Statistical Analysis

Wilcoxon signed-rank tests were conducted to determine statistically significant crossmodal congruency effects (differences in means between congruent and incongruent trials) for each participant. Only participants with significant CCE scores were included in comparisons across modalities for both the knee and foot. A linear Pearson’s correlation analysis was performed to determine any association between participant’s naturalness ratings and crossmodal congruency effects of each modality. For group-level analysis, a Wilcoxon signed rank test was performed to determine any differences in CCE scores or in number of incorrect responses. For visual reaction times, a one-way analysis of variance followed by post-hoc multiple comparison test was conducted to determine significant differences between reaction times for the visual stimulus at the knee or the foot. For the EEG band power analysis, we used Wilcoxon signed-rank tests to determine significant differences between the incongruent and congruent conditions for each electrode, followed by false discovery rate correction. For the group-level analysis, we used Wilcoxon rank-sum test. An alpha of 0.05 was used for all statistical analyses.

## References

1. Viseux, F. J. F. The sensory role of the sole of the foot: Review and update on clinical perspectives. Neurophysiol. Clin. 50, 55–68 (2020).

2. Lundborg, G. & Rosen, B. SENSORY SUBSTITUTION IN PROSTHETICS. Hand Clin. 17, 481–488 (2001).

3. Raspopovic, S., Valle, G. & Petrini, F. M. Sensory feedback for limb prostheses in amputees. Nat. Mater. 20, 925–939 (2021).

4. Petros, E. et al. Long-term performance and stability of implanted neural interfaces in individuals with lower limb loss. J. Neural Eng. 22, 016013 (2025).

5. Nanivadekar, A. C. et al. Restoration of sensory feedback from the foot and reduction of phantom limb pain via closed-loop spinal cord stimulation. *Nat*. Biomed. Eng. 1–12 (2023) doi:10.1038/s41551-023-01153-8.

6. Kristjánsson, Á. et al. Designing sensory-substitution devices: Principles, pitfalls and potential1. Restor. Neurol. Neurosci. 34, 769–787.

7. Heller, B. W., Datta, D. & Howitt, J. A pilot study comparing the cognitive demand of walking for transfemoral amputees using the Intelligent Prosthesis with that using conventionally damped knees. Clin. Rehabil. 14, 518–522 (2000).

8. Sensinger, J. W. & Dosen, S. A Review of Sensory Feedback in Upper-Limb Prostheses From the Perspective of Human Motor Control. Front. Neurosci. 14, (2020).

9. Tan, D. W. et al. A neural interface provides long-term stable natural touch perception. Sci. Transl. Med. 6, (2014).

10. Saal, H. P. & Bensmaia, S. J. Biomimetic approaches to bionic touch through a peripheral nerve interface. Neuropsychologia 79, 344–353 (2015).

11. George, J. A. et al. Biomimetic sensory feedback through peripheral nerve stimulation improves dexterous use of a bionic hand. *Sci*. Robot. 4, eaax2352 (2019).

12. Okorokova, E. V., He, Q. & Bensmaia, S. J. Biomimetic encoding model for restoring touch in bionic hands through a nerve interface. J. Neural Eng. 15, 066033 (2018).

13. Valle, G. et al. Biomimetic Intraneural Sensory Feedback Enhances Sensation Naturalness, Tactile Sensitivity, and Manual Dexterity in a Bidirectional Prosthesis. Neuron 100, 37–45.e7 (2018).

14. Chandrasekaran, S. et al. Sensory restoration by epidural stimulation of the lateral spinal cord in upper-limb amputees. 10.7554/eLife.54349 doi:10.7554/eLife.54349.

15. Walker, P., Francis, B. J. & Walker, L. The Brightness-Weight Illusion. Exp. Psychol. 57, 462–469 (2010).

16. Cross-Modal Interaction between Vision and Touch: The Role of Synesthetic Correspondence - Gail Martino, Lawrence E Marks, 2000. https://journals.sagepub.com/doi/10.1068/p2984.

17. Blustein, D., Wilson, A. & Sensinger, J. Assessing the quality of supplementary sensory feedback using the crossmodal congruency task. Sci. Rep. 8, (2018).

18. Spence, C. Crossmodal correspondences: A tutorial review. Atten. Percept. Psychophys. 73, 971–995 (2011).

19. Haciahmet, C. C., Frings, C., Beste, C., Münchau, A. & Pastötter, B. Posterior delta/theta EEG activity as an early signal of Stroop conflict detection. Psychophysiology 60, e14195 (2023).

20. Forster, B. & Pavone, E. F. Electrophysiological correlates of crossmodal visual distractor congruency effects: Evidence for response conflict. Cogn. Affect. Behav. Neurosci. 8, 65–73 (2008).

21. Kanayama, N., Sato, A. & Ohira, H. The role of gamma band oscillations and synchrony on rubber hand illusion and crossmodal integration. Brain Cogn. 69, 19–29 (2009).

22. Marasco, P. D. et al. Neurorobotic fusion of prosthetic touch, kinesthesia, and movement in bionic upper limbs promotes intrinsic brain behaviors. *Sci*. Robot. 6, eabf3368 (2021).

23. Petrini, F. M. et al. Sensory feedback restoration in leg amputees improves walking speed, metabolic cost and phantom pain. Nat. Med. 25, 1356–1363 (2019).

24. Charkhkar, H. et al. High-density peripheral nerve cuffs restore natural sensation to individuals with lower-limb amputations. J. Neural Eng. 15, 056002 (2018).

25. Zopf, R., Savage, G. & Williams, M. A. Crossmodal congruency measures of lateral distance effects on the rubber hand illusion. Neuropsychologia 48, 713–725 (2010).

26. Spence, C., Pavani, F., Maravita, A. & Holmes, N. Multisensory contributions to the 3-D representation of visuotactile peripersonal space in humans: Evidence from the crossmodal congruency task. J. Physiol. Paris 98, 171–189 (2004).

27. Occelli, V., Spence, C. & Zampini, M. The effect of sound intensity on the audiotactile crossmodal dynamic capture effect. Exp. Brain Res. 193, 409–419 (2009).

28. Klöppel, S. et al. The effect of handedness on cortical motor activation during simple bilateral movements. NeuroImage 34, 274–280 (2007).

29. Shen, Y.-C. & franz, E. A. Hemispheric Competition in Left-Handers on Bimanual Reaction Time Tasks. J. Mot. Behav. 37, 3–9 (2005).

30. Corniani, G. & Saal, H. P. Tactile innervation densities across the whole body. J. Neurophysiol. 124, 1229–1240 (2020).

31. Igarashi, Y., Kimura, Y., Spence, C. & Ichihara, S. The selective effect of the image of a hand on visuotactile interactions as assessed by performance on the crossmodal congruency task. Exp. Brain Res. 184, 31–38 (2008).

32. Spence, C., Nicholls, M. E. R., Gillespie, N. & Driver, J. Cross-modal links in exogenous covert spatial orienting between touch, audition, and vision. Percept. Psychophys. 60, 544– 557 (1998).

33. Spence, C., Pavani, F. & Driver, J. Spatial constraints on visual-tactile cross-modal distractor congruency effects. Cogn. Affect. Behav. Neurosci. 4, 148–169 (2004).

34. Marini, F., Romano, D. & Maravita, A. The contribution of response conflict, multisensory integration, and body-mediated attention to the crossmodal congruency effect. Exp. Brain Res. 235, 873–887 (2017).

35. Hutchison, B. C. et al. Perceptual Differences between Cortical and Peripheral Stimulation Strategies for Sensory Restoration. 2025.08.20.25334094 Preprint at 10.1101/2025.08.20.25334094 (2025).

36. Mancini, F. et al. Whole-body mapping of spatial acuity for pain and touch. Ann. Neurol. 75, 917–924 (2014).

37. Le Bars, D. The whole body receptive field of dorsal horn multireceptive neurones. Brain Res. Rev. 40, 29–44 (2002).

38. Blustein, D., Gill, S., Wilson, A. & Sensinger, J. Crossmodal congruency effect scores decrease with repeat test exposure. PeerJ 7, e6976 (2019).

39. Holmes, N. P., Calvert, G. A. & Spence, C. Tool use changes multisensory interactions in seconds: evidence from the crossmodal congruency task. Exp. Brain Res. 183, 465–476 (2007).

40. Poole, D., Couth, S., Gowen, E., Warren, P. A. & Poliakoff, E. Adapting the Crossmodal Congruency Task for Measuring the Limits of Visual–Tactile Interactions Within and Between Groups. Multisensory Res. 28, 227–244 (2015).

41. Sengül, A. et al. Extending the Body to Virtual Tools Using a Robotic Surgical Interface: Evidence from the Crossmodal Congruency Task. PLOS ONE 7, e49473 (2012).

42. Nebes, R. D. Vocal Versus Manual Response As a Determinant of Age Difference in Simple Reaction Time1. J. Gerontol. 33, 884–889 (1978).

43. Pastötter, B. & Frings, C. It’s the Other Way Around! Early Modulation of Sensory Distractor Processing Induced by Late Response Conflict. J. Cogn. Neurosci. 30, 985–998 (2018).

44. Marly, A., Yazdjian, A. & Soto-Faraco, S. The role of conflict processing in multisensory perception: behavioural and electroencephalography evidence. Philos. Trans. R. Soc. B Biol. Sci. 378, 20220346 (2023).

45. Cohen, M. X. & Ridderinkhof, K. R. EEG Source Reconstruction Reveals Frontal-Parietal Dynamics of Spatial Conflict Processing. PLOS ONE 8, e57293 (2013).

46. Harmony, T. The functional significance of delta oscillations in cognitive processing. Front. Integr. Neurosci. 7, (2013).

47. Karakaş, S. A review of theta oscillation and its functional correlates. Int. J. Psychophysiol. 157, 82–99 (2020).

48. H, J. H. Ten-twenty electrode system of the international federation. Electroencephalogr Clin Neurophysiol 10, 371–375 (1958).

49. Bose, R. et al. Classification of brain signal (EEG) induced by shape-analogous letter perception. Adv. Eng. Inform. 42, 100992 (2019).

50. Abbasi, N. I., Bose, R., Bezerianos, A., Thakor, N. V. & Dragomir, A. EEG-Based Classification of Olfactory Response to Pleasant Stimuli. in Proceedings of the Annual International Conference of the IEEE Engineering in Medicine and Biology Society, EMBS (2019). doi:10.1109/EMBC.2019.8857673.

