## Supplemental Material for "The Crossmodal Congruency Task as a measure of intuitiveness of sensory feedback in the lower limb"

### Crossmodal Congruency Task as a measure of intuitiveness of sensory feedback – A feasibility study

#### Visual Reaction Time

*Supplementary Table 1: P-values of One-way ANOVA of visual reaction times across two location and three luminous intensity*

| Subject | P-value |
|---|---|
| 1 | 0.37 |
| 2 | $0.12 \times 10^{-5}$ |
| 3 | 0.02 |
| 4 | 0.26 |
| 5 | 0.85 |
| 6 | 0.51 |
| 7 | 0.77 |
| 8 | 0.76 |
| 9 | 0.005 |
| 10 | 0.14 |
| 11 | 0.82 |
| 12 | 0.44 |
| 13 | 0.28 |
| 14 | 0.23 |
| 15 | 0.15 |

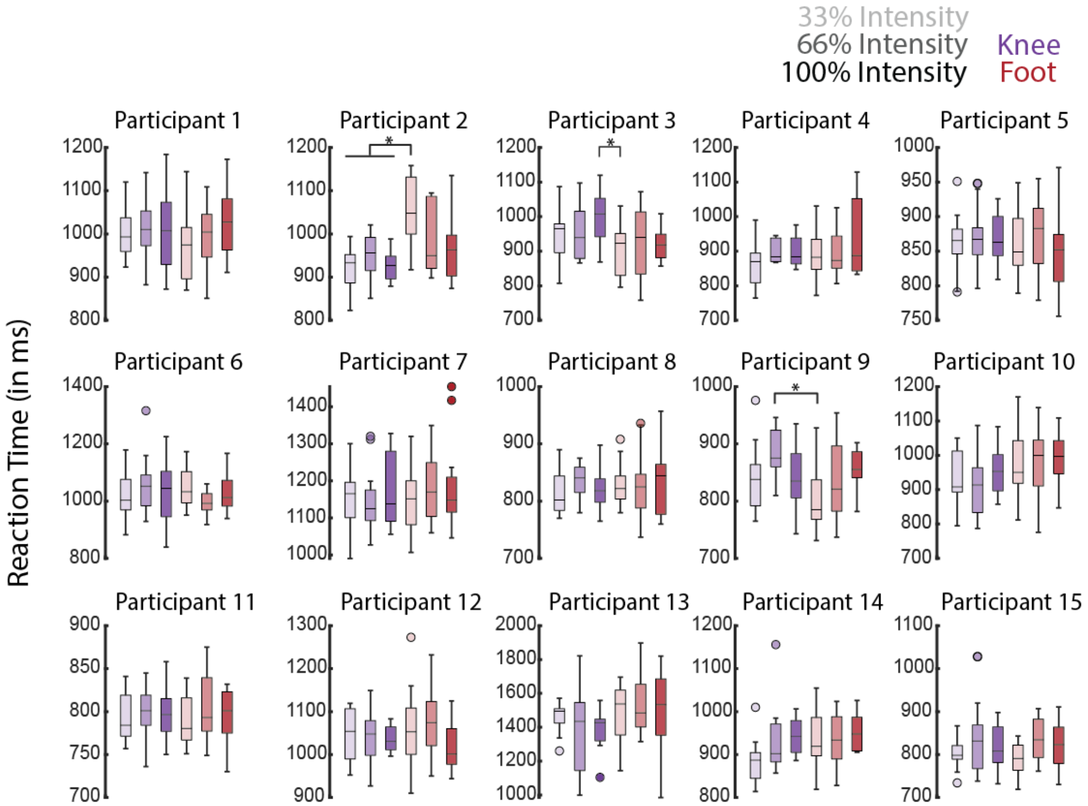


**Supplementary Fig 1:** Reaction time for visual stimuli only for each participant at the knee (purple) and foot (red). Lighter to darker shades indicate the luminous intensity of the stimulus. The * indicates significant difference (p<0.05) based on the post-hoc multiple comparisons after one-way ANOVA.

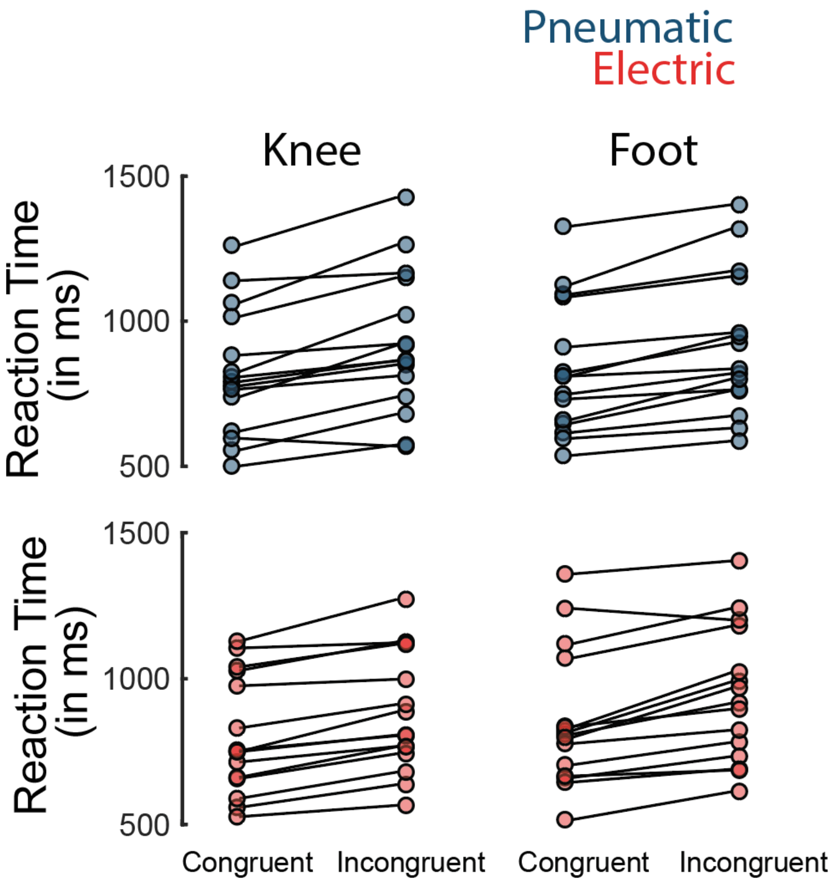


**Supplementary Fig 2:** Reaction time for congruent and incongruent trials for each participant in the CCE task. Top row shows the reaction time for the pneumatic (blue) stimulus and the bottom row represents the reaction time for the electric (red) stimulus.

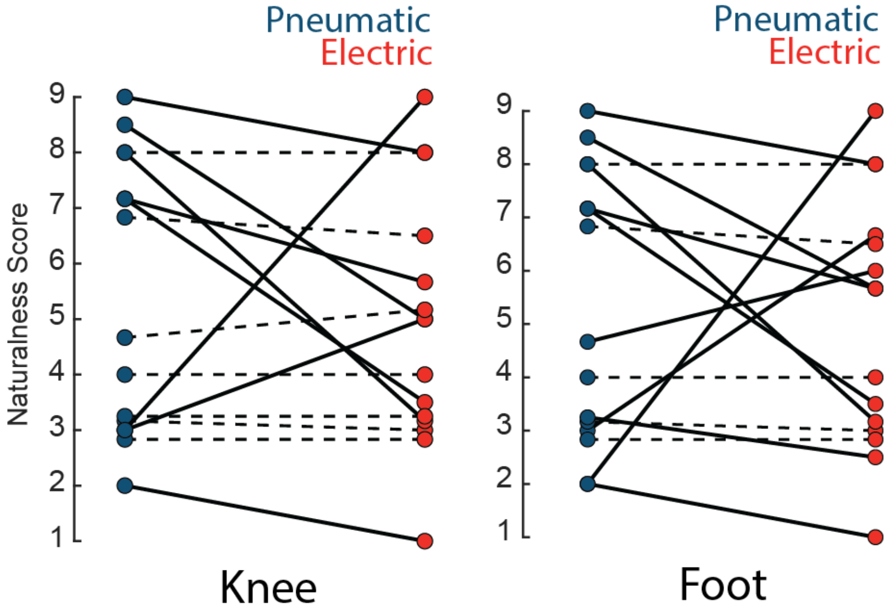


**Supplementary Fig 3:** Subjective naturalness rating for the pneumatic (blue) and electric (red) stimulus at the knee and foot. The solid lines represent significant difference between the Pneumatic and Electric stimulus ($p<0.05$, bootstrapping) and dashed lines represent no significant difference.

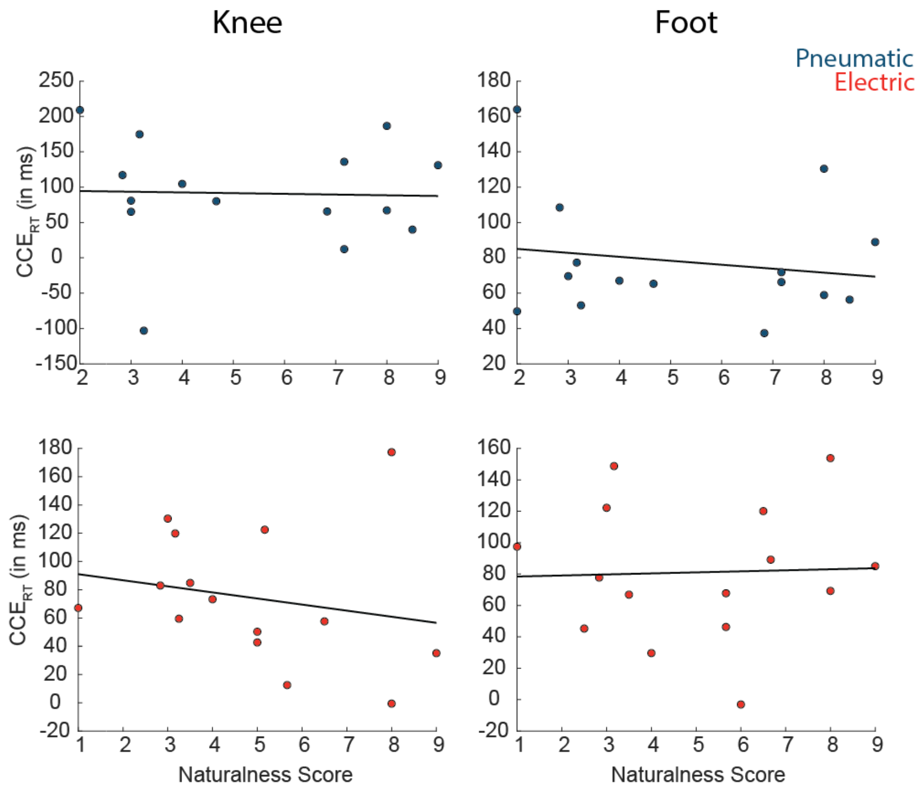


**Supplementary Fig 4:** Correlation across participants between naturalness and CCE scores for pneumatic (A,B) and electric (C,D) stimulus for the knee and foot. We did not observe a significant inter-participant correlation (Pearson's r) between the CCE scores and the perceived naturalness ratings for pneumatic-knee ($r=-0.03$, $p=0.91$), pneumatic-foot ($r=-0.17$, $p=0.54$), electric-knee ($r=-0.20$, $p=0.47$) or electric-foot ($r=0.04$, $p=0.9$) conditions.

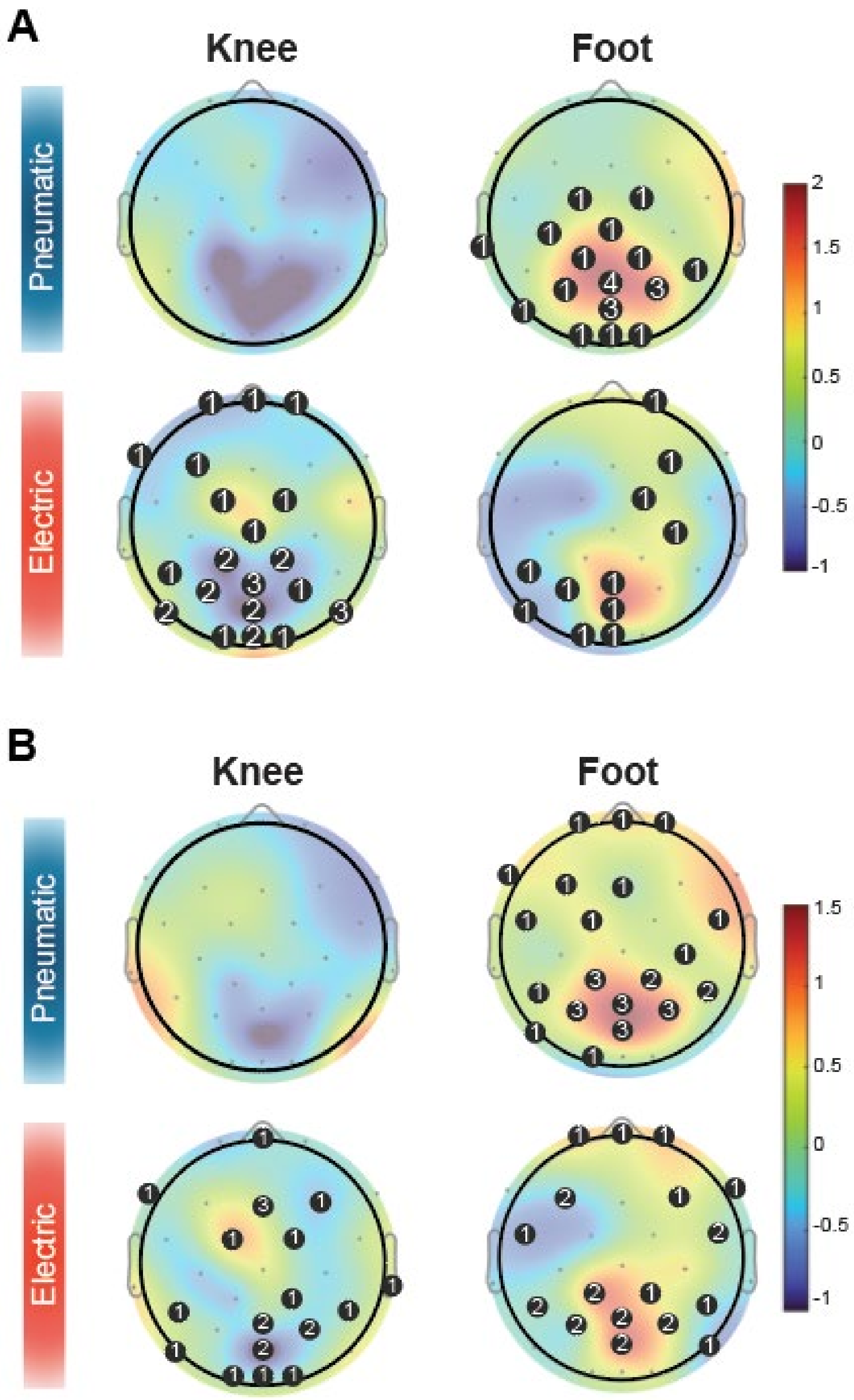


**Supplementary Fig 5:** Stim-locked (A) delta and (B) theta band power results. (A) The topographic plots show the difference in the incongruent and congruent conditions for pneumatic (blue) and electric (red) stimuli. The black circles indicate the number of participants that had a significant difference in that electrode.

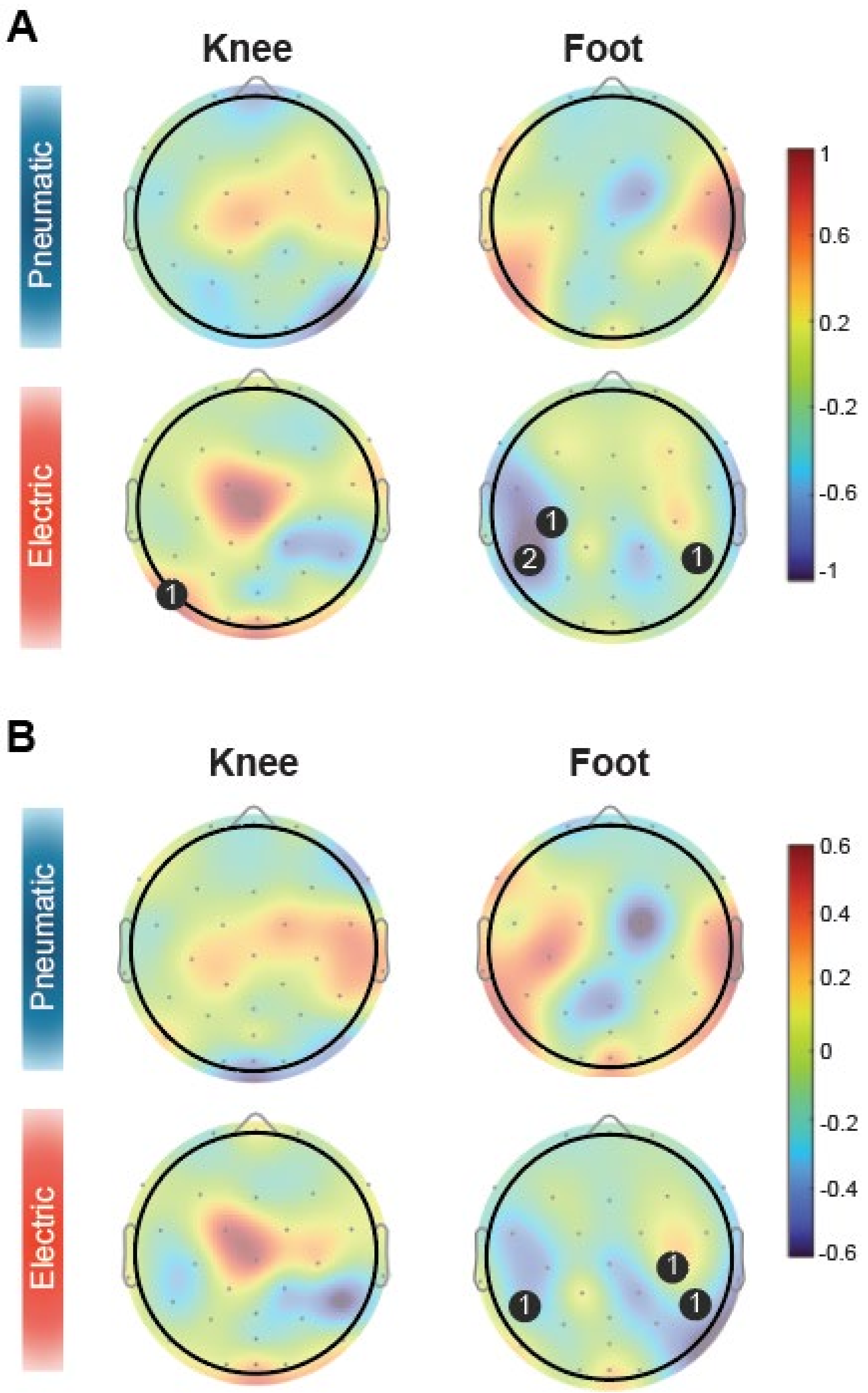


**Supplementary Fig 6:** Response-locked (A) delta and (B) theta band power results. (A) The topographic plots show the difference in the incongruent and congruent conditions for pneumatic (blue) and electric (red) stimuli. The black circles indicate the number of participants with a significant difference at that electrode.

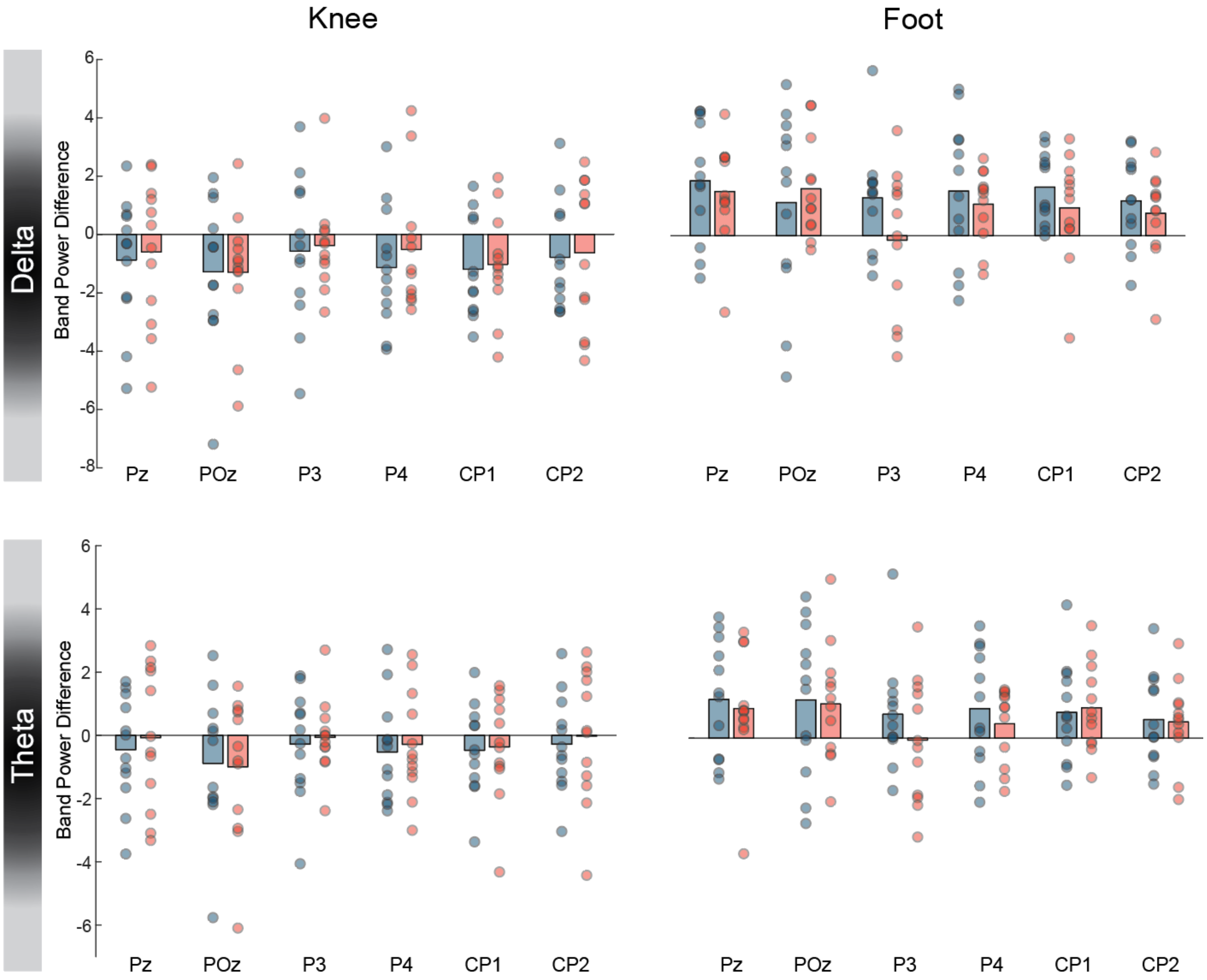


**Supplementary Fig 7:** Mean band power difference of the parietal electrodes between incongruent and congruent conditions for delta and theta band at the knee and foot locations. No significant difference was observed across any of the electrodes.

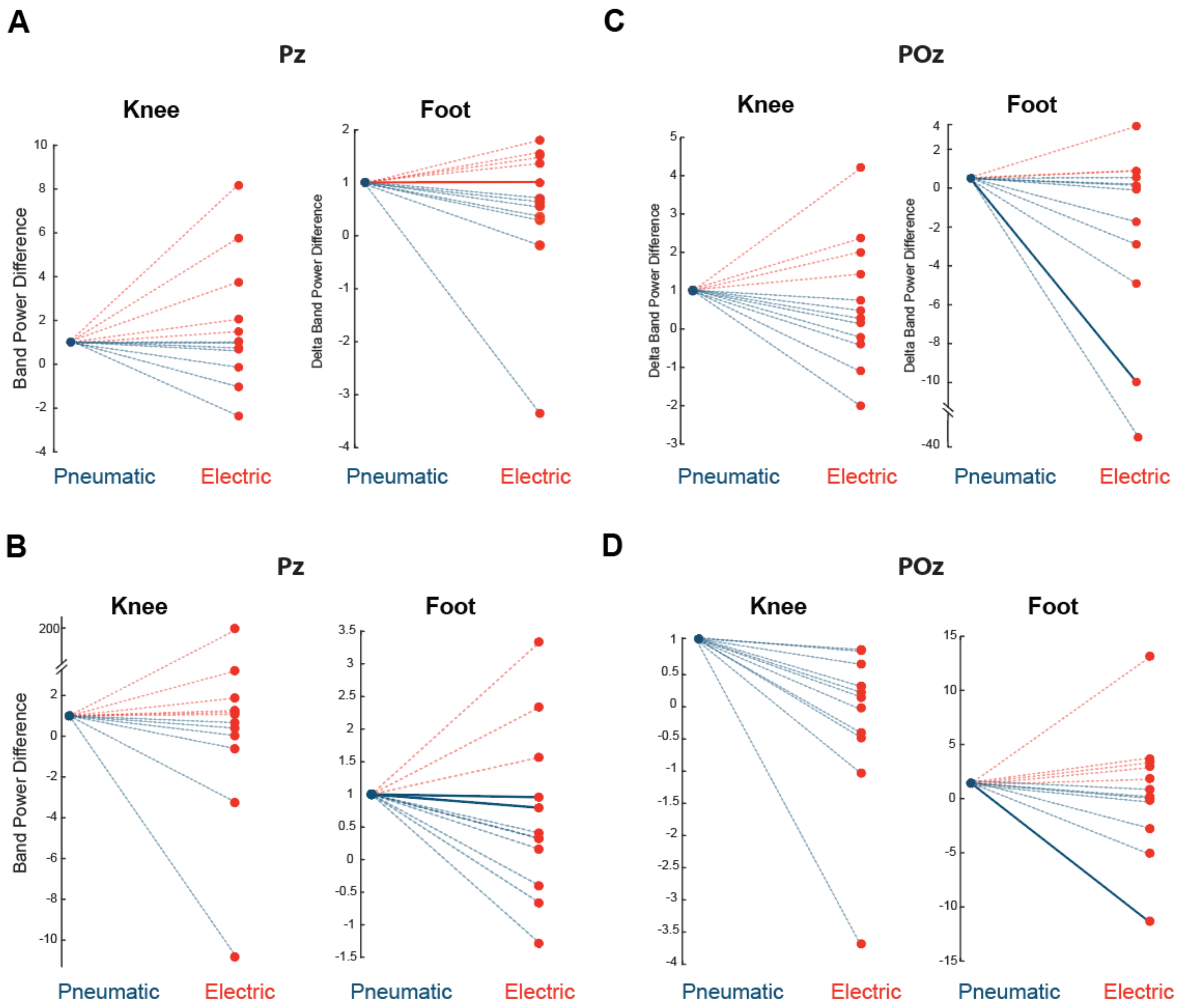


**Supplementary Fig 8:** Difference in the (A) delta and (B) theta band power for the Pz and POz electrodes between the incongruent and congruent conditions for the electric stimulus normalized to the pneumatic stimulus for each participant. The solid lines indicate a significant difference between incongruent and congruent conditions ($p<0.05$) for both pneumatic and electrical stimulus for that participant.
